# Breast Cancer Reaction-Diffusion from Spectral-Spatial Analysis in Immunohistochemistry

**DOI:** 10.1101/2022.07.10.499460

**Authors:** Stefano Pasetto, Mohammad U. Zahid, Roberto Diaz, Michael Montejo, Marilin Rosa, Robert Gatenby, Heiko Enderling

## Abstract

Cancer is a prevalent disease, and while many significant advances have been made, the ability to accurately predict how an individual tumor will grow – and ultimately respond to therapy – remains limited. We use spatial-spectral analysis of 20 patients accrued to a phase II study of preoperative SABR with 9.5 x 3 Gy for early-stage breast cancer whose tissues were stained with multiplex immunofluorescence. We employ the reaction-diffusion framework to compare the data-deduced two-point correlation function and the corresponding spatial power spectral distribution with the theoretically predicted ones. A single histopathological slice suffices to characterize the reaction-diffusion equation dynamics through its power spectral density giving us an interpretative key in terms of infiltration and diffusion of cancer on a per-patient basis. This novel approach tackles model-parameter-inference problems for tumor infiltration and can immediately inform clinical treatments.

## 1 Introduction

Cancer is a polymorphic system that evolves over many scales, from intracellular through signaling pathways, macromolecular trafficking, intercellular through cell-cell adhesion and communication, and tissue level through cell-extracellular matrix interaction mechanical forces (Hatzikirou et al., 2012). The complex interaction and co-evolution of the tumor with the host immune system recently received significant attention from cancer therapies that boost the immune system’s antitumor response. Therefore, it is imperative to characterize tumor topology and heterogeneity and its overall growth and invasion potential in the complex tissue environment.

Great attention has been given to a mathematical formalization of growth and diffusion processes for cancer (Gatenby and Gawlinski 1996). The most significant mathematical advancement in using continuous diffusion models to study cancer dynamics comes from brain cancer modeling (Swanson et al., 2000, 2001, 2002, 2003). General studies of tissue invasion (hypotaxis) analyze wave solutions and the tracking of heat-shock proteins (Szymanska et al., 2009), or hypoxic cell waves and the pseudo palisades (hypercellular regions) in glioblastoma (GBM) (Martinez-Gonzalez et al., 2012) in connection with imaging (Wang et al., 2009). Similarly, breast cancer cell diffusion and infiltration have been demonstrated to morph the extracellular matrix architecture (Pally et al., 2019). Diffusion equations have mainly been used in compartment model approaches, e.g., for evolution from low to high-grade glioma (Bogdańska et al., 2017) and its response to radiotherapy fractionation (Henares-Molina et al., 2017). More recently, reaction-diffusion equations have been deployed on the cellular level to simulate spatial cell size distributions and their influence on uptake rates (Gerlee and Nelander 2012, Klank et al., 2018, Khetan et al., 2019).

In most of these works, reaction-diffusion models are assumed as instruments to interpret a clinical condition (e.g., the anamnesis of a patient, a longitudinal past collection of data, and similar) and forecast tumor dynamics with and without therapeutic interventions. A remarkable example is a study of DTI-MRI-derived brain biomechanics that can predict the location of secondary cancer foci distant to the primary tumor (Angeli et al., 2018).

The diffusion process models’ role in the approach to the clinically significant problems of patient-specific imaging-derived predictive models in response to neoadjuvant breast cancer therapy is no less critical. Several authors studied this problem (Weis et al., 2013, 2015, 2017, Roque et al., 2018, Jarrett et al., 2018, Mang et al., 2020), focusing on the coupling diffusion model to extracellular matrix stiffness, in connection with radiotherapy (Roniotis et al., 2012, Holdsworth et al., 2012, Borasi et al., 2016) or synergy of radiation therapy and chemotherapy (Kibis and Buyuktahtakin 2018), surgical resection (Hathout et al., 2016), necrosis density thresholds (Patel and Hathout 2017), radiation-induced necrosis (Narasimhan et al., 2019), in vitro treatment of triple-negative breast cancer cell lines (Bowers et al., 2020), in connection with in vitro growth (Ayensa-Jiménez et al., 2020) or synthetic models of solid tumors (Sego et al., 2020).

Parallel to this exploding interest in imaging, and thanks to the new or enhanced information from these new techniques, is the effort to develop new and more effective mathematical and statistical methodologies that can extract the correct information from this wealth of data. We present a technique that contributes to interpreting these data in this work. Nevertheless, the technique is generalizable, applicable to any imaging set, and limited neither to 2D images nor to cancer diseases. In what follows, we will focus on image analysis of stained histopathological slides sampled from breast cancer patients:

- We stained and analyzed tumor slides with multiplex immunofluorescence acquired from biopsy pre and post-SABR;
- We analyze the spatial cell distribution of patient histopathology-stained slides (Fig. 1A) with the 2-point correlation function;
- We computed the corresponding power spectrum of the spatial fluctuation;
- We derived a new analytical formula for the power spectrum of the fluctuation function of a reaction-diffusion equation;
- We solve the inference problem between the reaction-diffusion equation and the data power spectral distribution

**Figure 1.**
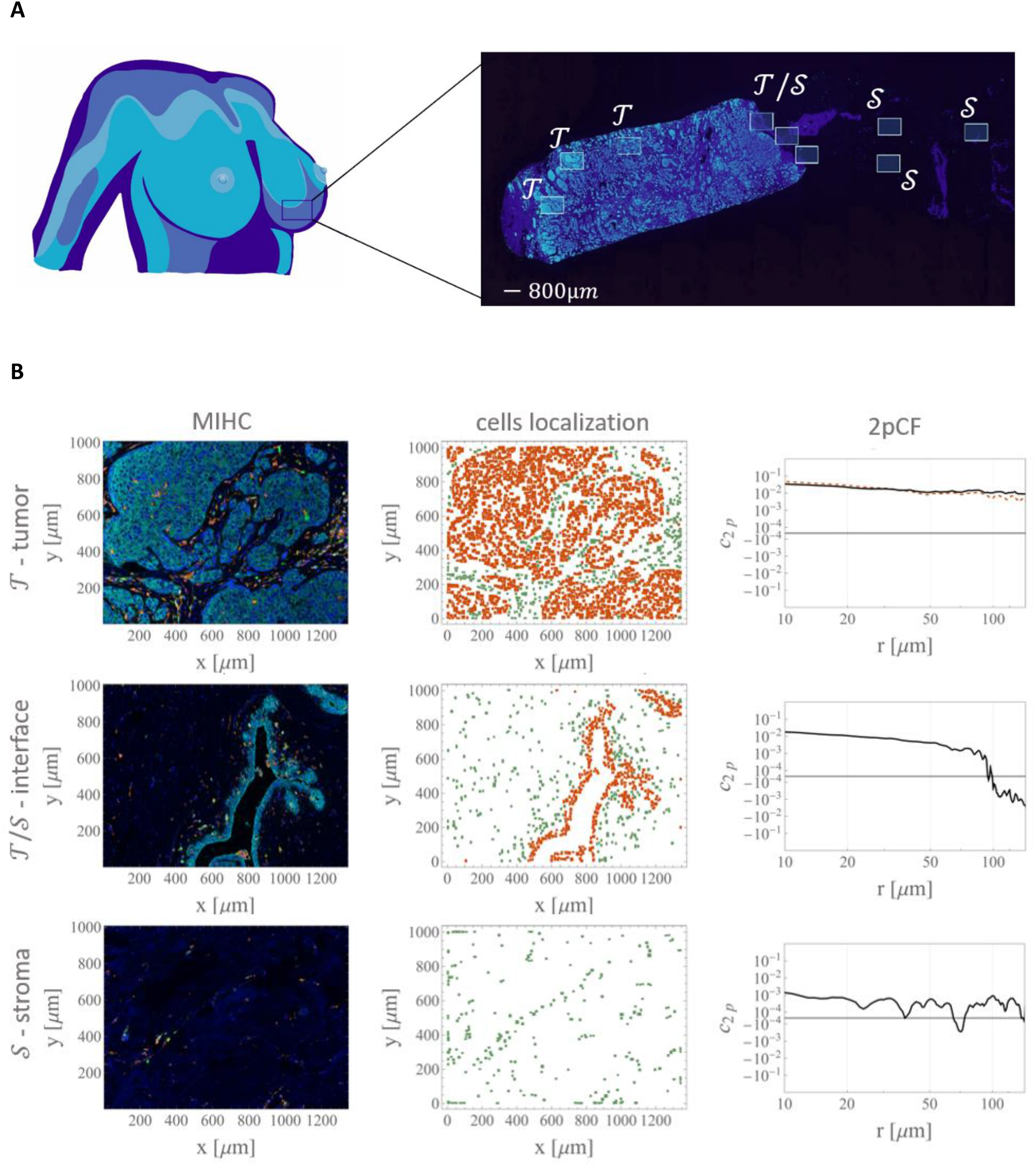
Patients’ resection area analysis. **Panel A**: a pictorial representation of the tissue resection areas process. **Panel B**: stained images (left column), cell localization (central column), and 2pCF (right column) for the tumoral, interface, and stromal ROI (top, central and bottom rows, respectively). The brown dashed curve in the top panel of the right column shows the 2pCF for only cancer cells. We use red for the tumor and green for all the other components in the localization plot column. In the third column, we see the characteristic trends of the 2pCF for ROI with the presence of cancer (solid, positive correlation), for the interface areas (dropping curve with anticorrelation in the outer ROI), and for the stroma ROI (small positive correlation with random irregular areas of anticorrelation) bordering the noise frequency detection thresholds.

This statistical tool introduced in the next section will show how cancer cell density varies as a function of the distance *r* from any cell (Riley et al., 2006). Three Regions of Interest (ROI) are selected, including tumor 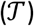, stroma distant to the tumor 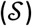, tumor and stroma interface (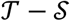, Fig. 1B).

## 2 Methods

### 2.1 Patient data

A total of *N*_pat_ = 20 patients have been accrued to a Phase II Study of preoperative SABR with three times 9.5 Gy for early-stage breast cancer (NCT03137693). A 5 μm thick unstained tumor slide from a formalin-fixed paraffin-embedded tumor block was acquired from biopsy and post-SABR and analyzed by the Moffitt Cancer Center tissue core and digital imaging laboratory. Tissues were stained with multiplex immunofluorescence to evaluate the infiltration of antigen-presenting cells, T-cells, NK cells, B-cells, and tumor cells using different combinations of CD3 and CD4 CD8, Foxp3, PD-1, CD68, PanCytoKeratin (PCK). For six patients, we have both pre-and post-treatment data. We focus on this subset of patients in the main text, leaving into Supplement 1 the investigation of patients with either pre-or post-treatment alone.

### 2.2 2-point correlation function

We implement the 2-point correlation function (2pCF) as a spatial statistic estimator for our histopathology tissue slices (Ridgway et al., 2006; Mosaliganti et al., 2009; Cooper et al., 2010, 2011; Binder & Simpson, 2015). The spatial n-point autocorrelation function *c*_np_ is defined as the probability of finding n cells at coordinates ***x***_1_, ***x***_2_,… ***x**_n_* in a suitably defined reference frame *S*(***O, x***) centered in ***O*** with ***x**_i_* = {*x*_1_, *x*_2_, *x*_3_}_*i*_ as spatial coordinates for the *i* = 1, 2, 3,… *n*. If we refer to the cell type *j, j* ∈ ***T*** with ***T*** set of possible cell types 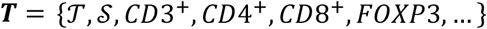, where 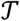 refers to tumor cell population, 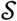-stroma cell population, and so forth. Then for the generic set type *j* in our histopathology-stained slide, we can formalize 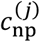 as 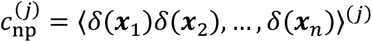(^1^). We focus on the 2pCF of the cancer population. We drop the index *j* and refer to the 2pCF as the auto-correlation function.

Furthermore, because of the connection we are going to develop in the next section with the continuous RD equation, we are going to implement for the 2pCF the following definition (^2^):

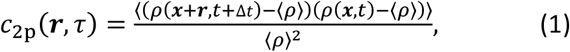

with ⟨*ρ*⟩ = *n* the number of cells for volume unit, and Δ*t* is a time interval. Because we work with a single breast cancer histopathology slice, we cannot sample the same tissue section twice, and we cannot consider the time dependence of the autocorrelation, and we will consider only ROI where large-scale homogeneity and isotropy of the cell distribution holds. Therefore we simplify Eq.(1) to:

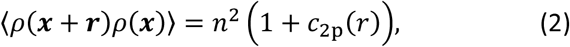

where *r* = ||**r**|| ≡ ||***x*_1_** – ***x***_2_||. The interpretation of this type of correlation function is simple: if the probability of finding a cell in the volume *dV* is *dP* = *ndV*, the average number of cells in the finite volume spanned by the histopathology slide is then ⟨*N*⟩ = *nV*, and the joined probability of finding two cells say cell 1 and cell 2, at a given distance *r*, is

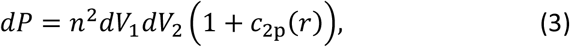

which also gives us a concept-interpretation key: If the cell distribution is a 3D random Poisson point process, the probability of finding cells in *dV*_1_ and *dV*_2_ are independent, in this case *c*_2p_ = 0, if the cell positions are correlated *c*_2p_ > 0, and if the positions are anticorrelated *c*_2p_ ∈ [−1,0[.

Note that to properly account for the 3D overlapping of cells in the spatial analysis of the 2pCF, we tested Eq. (2) with several kernels, boundary conditions, and bandwidth, always finding convergence (but not a coincidence) of results. We refer the interested reader to the technical treatment offered by the modulation-analysis presented in Supplement 2.

### 2.3 Power spectrum of the fluctuations

The Fourier transform of the autocorrelation function for a continuous function is the power spectrum. We only review the analogous for a 3D random point process we use in the following, focusing on and developing only the equations of interest (Alessio 2016). The Fourier transform for the distribution of cells with a density *ρ*(***x***, *t*) = (⟨*ρ*⟩*V*)^-1^ ∑_*i*_ *δ*(***x*** – ***x**_i_*(*t*)) is

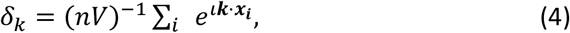

with wavenumber *k* = ||***k***||, with || || = || ||_2_ Euclidean norm, ***k*** = {***k_x_,k_y_,k_z_***}, where the barycenter of the *i*^th^ cell is located at ***x**_i_* and *e**^ιk·x_i_^*** is periodic in *V* (*ι* is the complex imaginary units, “·” the inner product). We slice the volume *V* (we refer to the slice and its volume with the same symbol *V* without loss of generality) into infinitesimal cells with unitary function **1**_*dv*_ (**1**_*dv*_ = 1 if the cell is in the volume *V*, or **1***_dv_* = 0 if it is not). Therefore Eq. (4) is equivalent to

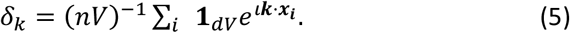

The averaged two-cells contribution to the spectrum of wavelengths, say cell 1 in *dV*_1_ and cell 2 in *dV*_2_, reads (note how because *δ*(**x**) is real the complex conjugate 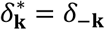):

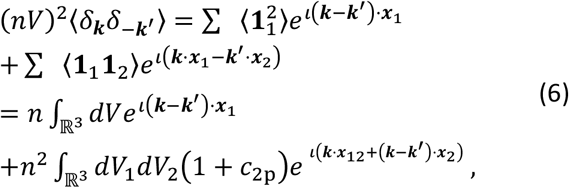

because ⟨**1**_1_**1**_2_⟩ = *n*^2^*dV*_1_*dV*_2_(1 + *c*_2p_) as results of considering Eq. (3) (with ***x***_12_ Euclidean distance between cell 1 and 2). Because Fourier components belonging to different ***k*** are statistically independent, the integrals in the previous vanishes if ***k* ≠ *k*^′^**, while ***k* = *k*^′^** yields, as a well-known result, the spectrum (the power spectral density, PSD) of the cell distribution:

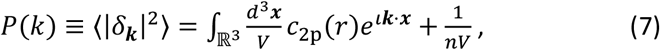

for ***k* ≠ 0**, and 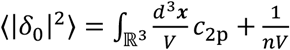 otherwise. The power spectrum measures the mean number of neighbors in excess of a random Poisson distribution within a distance of *~k*^-1^ from a randomly chosen cell. The extra term (*nV*)^-1^ is due to shot-noise (a contribution visible as white-noise *P*(*k*) ∝ *k*^0^). If the cell distribution is Poisson, then 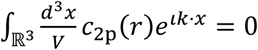, but the Fourier modes have a not null variance ⟨|*δ_k_*|^2^⟩ = (*nV*)^-1^ due to the discreteness. This classical result (Peebles 1973) is often coupled with filter-design windows functions commonly implemented in the power spectrum determination to optimize its analytical treatment (Dirichlet, Hamming, Blackman, etc.) that we also tested (see Supplement 2) with consistent results. In this paper, we will stick to the derivation presented above. Nuttall-softening length (Nuttall 1981) will be preferred if necessary.

### 2.4 2-point correlation function from the reaction-diffusion equation

Diffusion of a cluster of cells with density *ρ* = *ρ*(***r**, t*) at position ***r*** and at a time *t* is described by the combination of the continuity equation without source and sink terms as *∂_t_ρ* + ***∇. φ*** = 0 (with *∂* being the partial derivative and *∇* the gradient operator) and Fick’s first law of proportionality between flux ***φ*** and density gradient 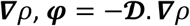, to get (for a generic anisotropic flow) 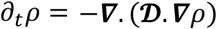 where 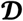 is the diffusion coefficient matrix. In a 3D system of reference (SoR) *S*(***O, r*** = {*x, y, z*}), with *r*^2^ = *x*^2^ + *y*^2^ + *z*^2^ being the squared radial direction from an arbitrary origin ***O***, we can write the diffusion equation for the locally homogeneous and isotropic case as 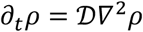, where *∇*^2^ is the D’Albertian operator. Green functions offer a natural solution for cell-like source points, even in the presence of an explicit exponential growth factor *γ*, i.e., 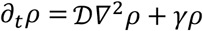 of interest here, in the form:

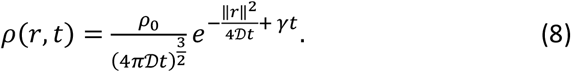

Note how this approach follows exponential tumor growth without carrying capacity constraints, and we will comment on this limitation later. For now, we focus on tumor RD statistics as an average over the tumor ROI, 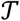. We assume that the cells are spread over a small 3D volume *d*^3^***r*** so that, as long as this volume is asymptotically smaller than the cell migration movement per unit time, i.e., to the first order in time, it holds on a linear scale 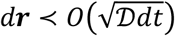, we can generalize the previous equation as 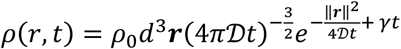. In the case of *n* different cells 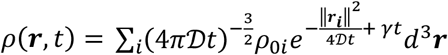 or, in the passage to the continuous limit, we get the classical heat-equation result that we are going to exploit here for our RD problem:

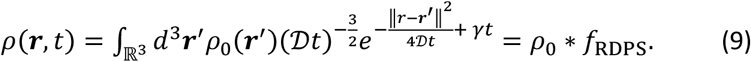

The last term in the equation is of interest here. The cell concentration at the generic time *t* is the convolution, denoted here by “*” of the initial concentration with a Gaussian, that by borrowing the terminology from Astronomy, we call RD point-spread-function *f*_RDPS_. Here cell concentration at +∞ is assumed to be null. Furthermore, by assuming constant temperature and pressure in the tissue sample, we justify 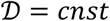. and elaborate on the *f*_RDPS_ as the function that carries the temporal dependence.

To proceed further, we now adopt a “fair representation” hypothesis: we assume that histopathology slides, surgically acquired from a local resection area, are reasonably representative of the clustering properties of the total tumor population of the patient. We comment more in the Discussion section on the implication of this assumption, which allows us to proceed with a mean-field solution for the RD equation over the histopathology slice volume. By performing this Gibbs average, we assume that the average volume formed is large enough to contain a sufficient number of cells to perform statistical analyses, but it is small enough to neglect large-scale gradients (i.e., 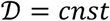 holds reasonably well). In this case, from Eq.(9) we obtain:

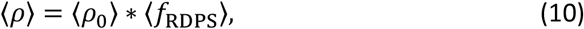

where we define the average over the time of the RD spread function as:

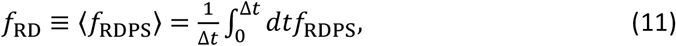

with Δ*t* sufficiently small that will indeed account for the non-equilibrium dynamics of diffusion and proliferation required by the average in *ρ*_0_. The drawback of this approach is that the same patient presents constant different diffusion coefficients 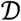 and growth rates *γ*, in distinct resection areas instead of having a single continuous function for 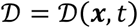 and the growth rate *γ* = *γ*(***x**, t*) dependent on spatial position. Furthermore, by visual inspection, we guess that the stochastic correlation is different from 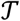 and 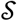 areas (Fig. 1), and we expect the theory to break down at the 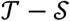 interface. Therefore, we are forced to solve the RD equation only locally in 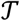 Finally, by considering the non-normalized autocorrelation of the average cell concentration ⟨*ρ*⟩, we write

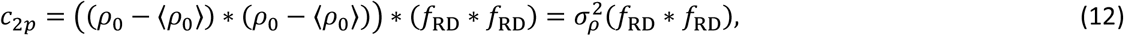

where the initial density is non-correlated, and the correlation function reduces to its variance *σ_ρ_*.The biological implication is that diffusing cells enhances cell proliferation – a visualization of loss of space and contact inhibition. Therefore, these reactions are stochastically correlated, and Eq. (12) assumes cell diffusion is the only mechanism by which cells become spatially correlated. The autocorrelation of *f*_RD_ captures this spatial correlation thus not contradicting Eq.(10) as ergodicity does not imply mixing considering our definition of averages (Supplement 3).

### 2.5 Power spectrum from reaction-diffusion equation

The RD equation’s analytical solution is unavailable for arbitrary reaction terms; therefore, it is easier to work with power spectral densities than with 2-point correlation functions to connect cancer diffusion with its spatial distribution. Performing Eq. (11) on Eq.(8), we obtain *f*_DR_ as

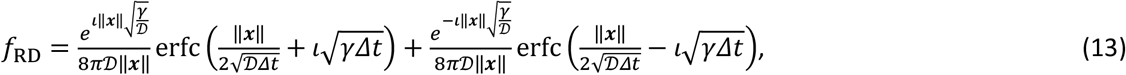

where erfc() is the complementary error function. Finally, considering Eq. (12), Eq.(13) together with the definition of power spectral density, we obtain

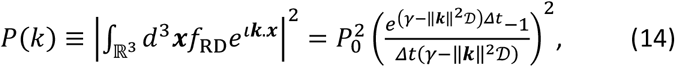

where for easier comparison with Eq. (7) we normalized with *P*_0_ to the zero-wavelength by computing 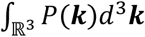 to obtain our final result.

We finally perform an inference exercise of the model Eq. (7) to the PSD data with the same Bayesian/Frequentists techniques (Pasetto et al., 2021a). Any other inference approach is equally valid.

## 3 Results

### 3.1 Breast tumor detection by the 2-point correlation function

Using the methodology developed in the previous section, we analyzed patient histopathology-stained spatial cell distribution (Fig. 1b). Qualitatively, we can recap the trend across the dataset.

The combined tumor and stroma cell distribution in a 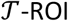 proposes a characteristic monomial distribution signature. Analysis of tumor cells’ spatial distribution follows a similar linear relationship in the bi-logarithmic plot. The correlation is mainly positive in both cases, meaning an architectural aggregation of the cells in the ROI exceeds the Poisson distribution (i.e., randomness). By contrast, cell clustering in stroma-dominated ROI, 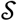, presents a more rapid loss of positive correlation to favor an irregularly oscillatory 2pCF profile at about one order of magnitude smaller values -cross patients-than for 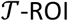. The 2pCF in 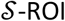 approximates *c*_2*p*_ ≅ 0, equivalent to a random cellular distribution, without any evident characteristic signature or regularity in the sample analyzed. Finally, the bi-logarithmic plot for intermediate 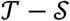 ROI presents a monomial behavior at low-medium *r*, similar to what is evidenced for 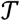 ROI. Still, it shows an abrupt drop that characterizes it as a significant negative *c*_2*p*_, i.e., at asymptotically larger *r* (less clustered than a random distribution).

### 3.2 Radiation therapy induces changes in cancer cell distribution morphology

We analyze the spatial distribution of cancer cells in the different 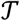 ROI before (Fig. 2A) and after (Fig. 2B) SABR. We investigate *c*_2*p*_ + 1 to evaluate only positive values of the 2pCF. The Poisson-process threshold moves from 0 to 1, with *c*_2_*_p_* > 1 representing spatial correlation or cell clustering, and *c*_2_*_p_* < 1 representing anticorrelation for 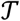 cell distribution alone. Cancer cells in three different 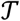 ROI, both pre-and post-treatment (referred to as A, B, and C in Fig 2 panel A, as well as D, E, and F in Fig. 2 panel B, respectively) strongly cluster in close neighborhoods (*r* < 50 μm) but lose positive correlation with a different characteristic slope as they approach Poisson randomness at more considerable distances (*r* > 50 μm; Fig. 2, the second column). Furthermore, we consider the 2pCF Fourier transform introduced in the previous section, the power spectral density (PSD), to better capture the multiscale contribution to the cancer architectural organization (tracked by wavenumbers, *k*) across the resection area. The characteristic “white-noise” plateau at small wavenumbers is followed after a characteristic wavenumber value where the PSD shape bends downward (sometimes referred to as correlation length) by the drop of the PSD at higher *k* (Fig. 2, third column).

**Figure 2.**
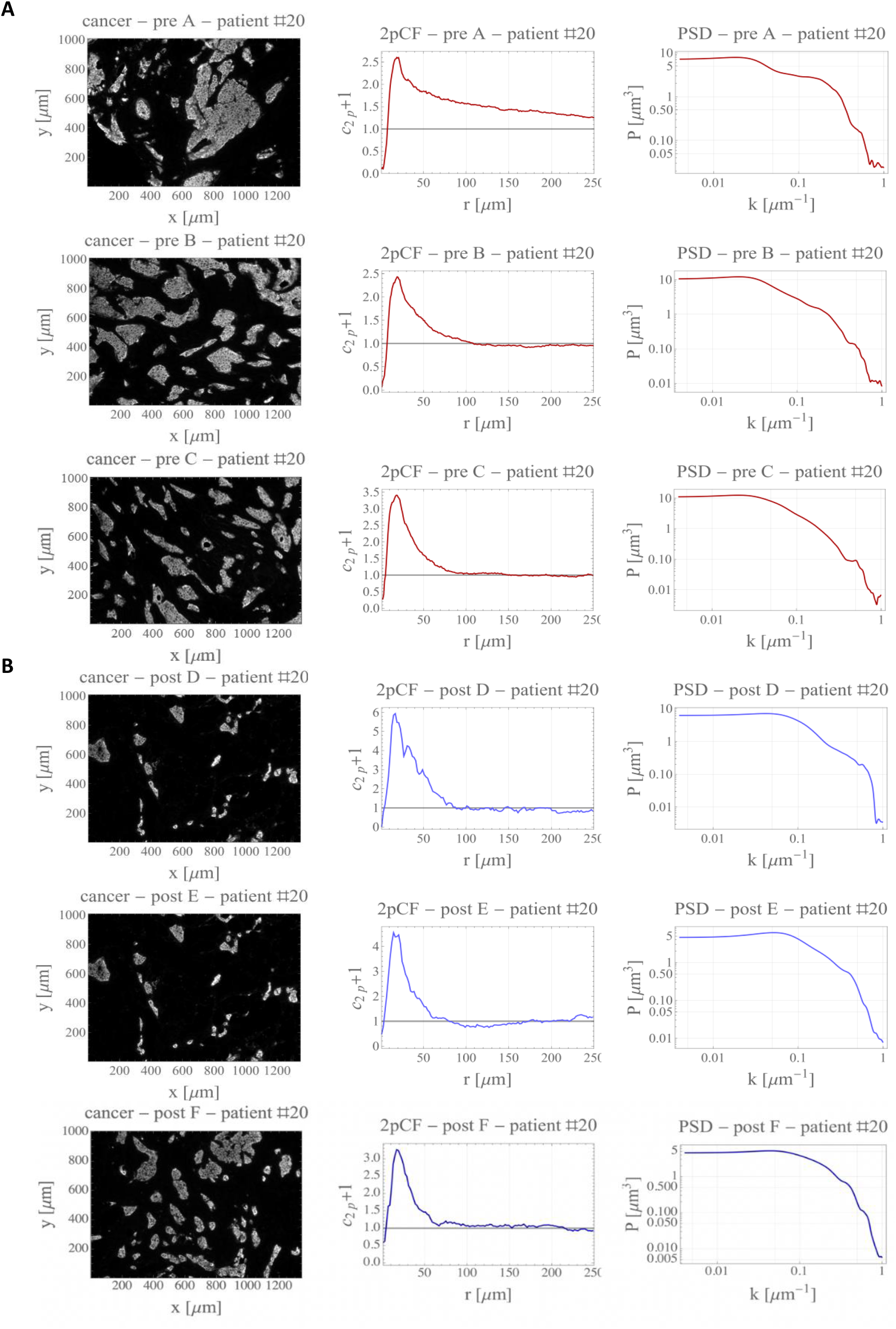
Power spectral analysis of cancer cell distribution. Panel a: (left column) spatial distribution monochromatic-stained for some patient-#20 tumoral resection areas (sites A, B, and C, the first three rows) referring to pre-treatment biopsy. On the central column, the corresponding 2pCF, on the right column the PSD. Panel B: as Panel A but for post-treatment biopsy.

### 3.3 The histopathology slides power spectrum analysis calibrates reaction-diffusion equation parameters

Patient tissues provide a single-time snapshot of the complex evolution of the cellular density in different spatial locations inside the resection area (Fig. 1). The exact location of each cancer cell and cancer cell conglomerates gives insights into the tumor proliferation and invasion dynamics. The reaction-diffusion (RD) equation represents a family of models relating the density of cancer cells in a given position at a given time as a function of the proliferation rate *γ* (per unit time) and a diffusion coefficient 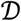 (area per unit time). We used the derived analytical formulation of the PSD for the homogeneous-isotropic RD equation and solved the RD parameter inference problem by interpreting the observed PSD of the histopathology slides with the PSD derived from the RD equation. We note how the RD framework disentangles the contributions between proliferation and diffusion at different wavenumbers, optimizing the inference solution. In Fig. 3a, we show how different reaction rates mainly contribute, and hence disentangle, at low wavenumbers while the influence at high *k* is dominated by the diffusion coefficient. The PSD of the RD equation closely captures the PSD of the pre-and post-treatment tissues (coefficient of determination *R*^2^ > 0.9; Fig. 3b).

**Figure 3.**
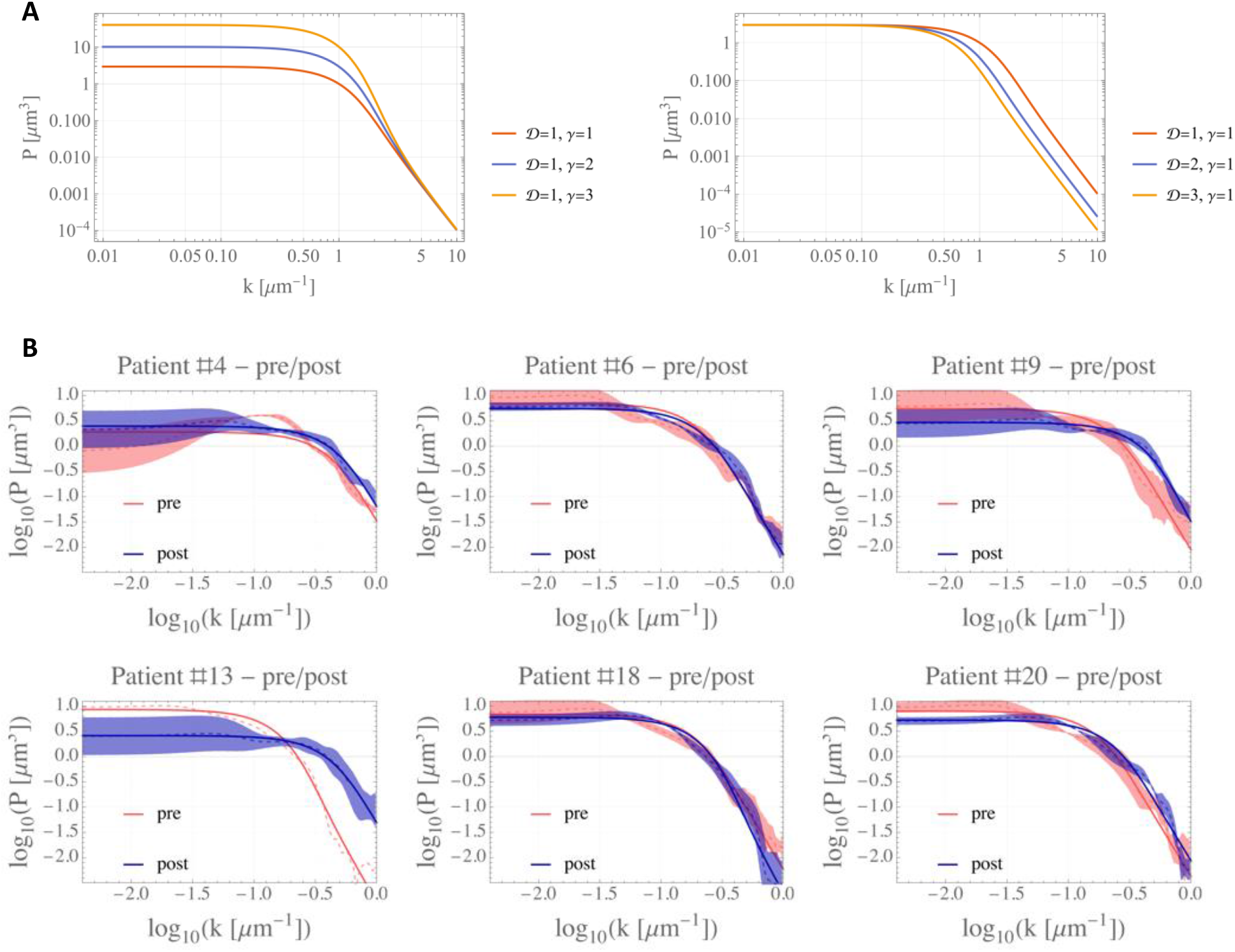
A. Power spectral density from RD framework. PSD contribution is shown for an arbitrarily chosen set of parameters to evidence the different contributions of the reaction and diffusion coefficients at different wavenumbers. B. Power spectral density pre-and post-treatment for several patients. The solid curve refers to the RD model fit of the three pre-SBRT ROI (A, B, C; red) and the three post-SPRT ROI (D, E, F; blue). Dashed curves are the averages, and transparent shaded areas show standard error.

Furthermore, the RD framework disentangles the different contributions between proliferation and diffusion by systematic differences in pre and post-treatment PSD. A direct comparison of pre-and-post-SABR RD parameters demonstrates a reduction in the patient-specific diffusion parameter, 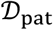 for each patient normalized by the average patient-specific diffusion parameter, 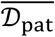 (i.e., across all 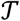 ROI pre and post-SABR, Fig. 4A). However, the normalized patient-specific growth rate parameter showed a dichotomy of responses for the 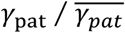 trend: increasing for some and decreasing for other patients (Fig. 4B). In Fig. 4C, the ratio of the coefficients (not averaged) is shown to track the cumulative effect of disease diffusion and proliferation: SBRT may induce an increase in the ratio with expected tumor growth (patients #6, #9, #18). In contrast, a decreasing trend suggests likely post-SBRT tumor regression (patients #4, #13, #20).

**Figure 4.**
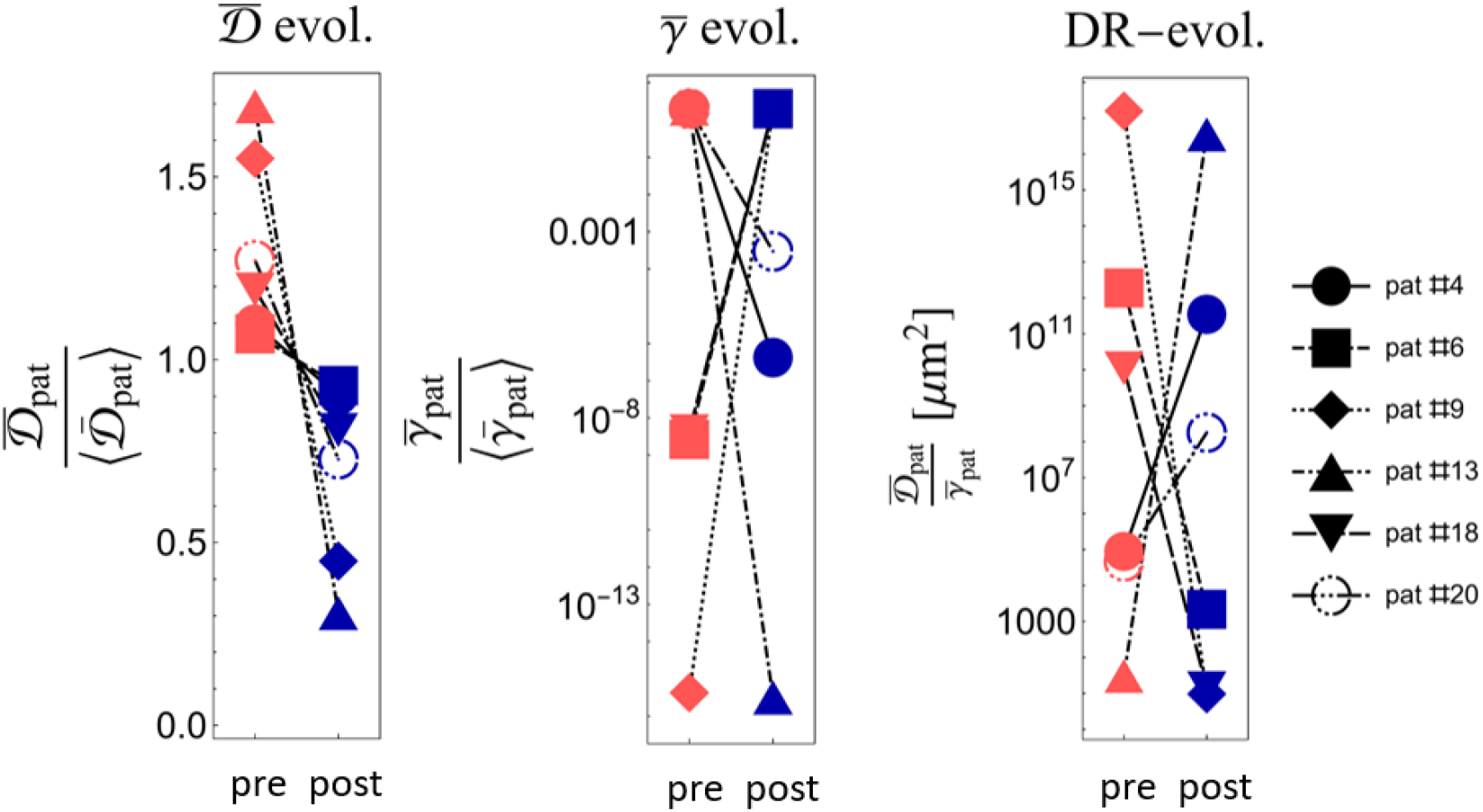
Comparison of pre-and-post-SABR reaction-diffusion parameters. **A.** Spatially averaged across all 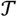 ROI per-patient diffusion parameters, 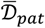, normalized by the time-averaged patient-specific diffusion parameter, 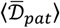, across all 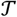 ROI for all times (pre and post). **B.** Same as A for the growth rates. **C.** Ration of the two coefficients.

## 4 Discussion

Cancer is a complex adaptive dynamic system that evolves at multiple scales; therefore, a trans-levels technique that captures the characteristics of several scales (or equivalently over several wavenumbers) is essential. Here we presented the first step in developing a novel interpretative methodology to capture the dominant scale lengths that drive breast cancer architectures before and after stereotactic ablative radiotherapy. Herein, we explore the potential of the cancer-cell architecture-defragmentation analysis and interpret it as a growth and tissue invasion process through the RD equation. The two-point correlation function connects spatial architecture and tumor growth and diffusion. The power spectral density is obtained directly from patient tissue immunofluorescence histopathology slides to calibrate the parameters of the RD equation.

Such an approach requires a few theoretical and biological hypotheses, particularly the “fair representation” hypothesis that histopathology slides are reasonably representative of the clustering properties of the total tumor population. While the tumor’s loco-regional properties may be well captured in an individual drop, the overall tumor is highly heterogeneous, and the cancer cell population’s local characteristics might depend on the micro-environment. We assumed that the sampled ROI is representative of the overall patient disease. The fair-representation hypothesis then permits us to generalize the consideration by comparing different patient biopsies, empowering our considerations. To achieve our goals in the present work, we took the validity of histopathology spatial characterization worked out with the 2pCF as fairly representing the nature of the patient breast cancer. While potentially conceived as a limitation; instead, this approach offers the possibility to control on a cell-by-cell basis the connection between spatial statistical tools, the two-point correlation function, and the corresponding power spectrum.

The second strong hypothesis is found in the choice of the interpretative framework: the RD equation. While the 2pCF and the corresponding PSD from the dataset are virtually free of assumption, the RD framework was not derived from the data but was chosen due to its simplicity. No reason a priori justifies the choice and form of the equation if not the practical analytical tractability, especially in Fourier space. While more physically involved approaches exist, such as Navier Stokes approaches, especially in the glioma literature (see introductive references), numerical investigation of integrative instruments and the power spectral density (which easily captures several orders of magnitude in wavelength) would involve multiscale physical parametrization that is not justified by the resolution and quantity of the available data. Thus, the chosen approach aligns with the clinical dataset and moves toward a predictive quantitative oncology framework. The clinical data set only provided matched pre-and-post-SBRT tissue samples for six patients with comparable outcomes for quantitative analysis. Therefore, this work focuses on developing breast cancer patient-specific reaction-diffusion dynamics from spectral analysis of immunohistochemistry slides. Future work will include analysis of such derived dynamics to predict complete pathological responses, relapse-free, and overall survival to help guide clinical decision-making (Pasetto et al., 2021b).

Finally, although beyond the scope of this study, it is conceivable that 2pCF analyses could help build a robust classifier to detect the presence of cancer architectures in selected biopsy ROI. Future work will include rigorous analysis of such digital pathology approaches.

## 5 Conclusion

Understanding tumor growth and invasion dynamics are paramount to personalizing patient care and improving outcomes. We developed a novel quantitative approach to infer such dynamic characteristics from a single biopsy taken at patient diagnosis. Inspired by looking at Fig. 4, we can easily envisage that for a patient described by a high proliferative index and a low diffusivity, we need to act with a targeted local aggressive therapy (e.g., radiation therapy). Vice versa, we want to proceed with systemic treatments (e.g., immunotherapy, chemotherapy) for a slowly proliferating cancer but highly diffusive and prone to metastasis. This result represents a fundamental step toward integrating quantitative methods into clinical decision-making to improve treatment responses and outcomes (Pasetto et al., 2021b). Future steps will require the development of a valid medical actionable path and extending the presented pilot project to a dataset with more substantial statistical significance. From one side, we will need further validation of the approach used, eventually to other cancer types, to understand the novel methodology’s range and limits of applicability. From the other, we will work to expand the statistical sample to make quantitative the forecasting ideas proposed here in order to transform the simple indications of Fig. 4 into a robust instrument able to predict the cancer evolution with achieved accuracy statistically.

## Acknowledgments

The authors would like to thank Isha Harshe for her help with Fig. 1. Funding: This work was supported by NIH/NCI grant U01CA244100; the Florida Breast Cancer Foundation; the JAYNE KOSKINAS TED GIOVANIS FOUNDATION FOR HEALTH AND POLICY, a private foundation committed to critical funding of cancer research. The opinions, findings, conclusions, or recommendations expressed in this material are those of the author(s) and not necessarily those of the JAYNE KOSKINAS TED GIOVANIS FOUNDATION FOR HEALTH AND POLICY, or their respective directors, officers, or staffs.

## Ethical statement

This study met the approval of the institutional review board, protocol MCC 18383: “Treatment Outcomes of Breast Cancer Patients Treated with All or One of the Multimodality Therapies at H. Lee Moffitt Cancer Center & Research Institute.”

## Supplement sections

### Supplement 1: Extended patient dataset

Of the 20 patients analyzed, ten presented clear tumor ROI (see Fig.1 main text), and between them, only six presented pre-and post-treatment biopsies. Nevertheless, the technique developed can infer the diffusion coefficient and reaction rate from a single timestep analysis of the 10 ROI. The extended main text Fig. 4c is in Fig. S2. No literature on 3D diffusion coefficients and reaction rates derived from single-cell resolution are available to date; therefore, pure coefficient values are not of interest here.

### Supplement 2: Modulation analysis

We can normalize the 2pCF to the unitary sphere in 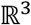 to test the impact of the most common non-parametric kernel function *κ*, wherefrom signal-analysis literature, we tested the uniform (default in this work) 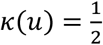, triangular *κ*(*u*) = 1 – |*u*|, parabolic 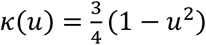 with support |*u*| ≤ 1, quadratic (biweight) 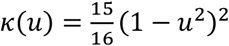 over |*u*| ≤ 1, cosine 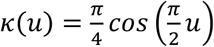 with support |*u*| ≤ 1 Gaussian, SemiCircle, and Triweight 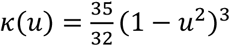 over |*u*| ≤ 1 (Epanechnikov 1969). Although many powerful parametric kernels might help the fitting process, we avoid arbitrary parameter introductions. By following the scheme in (Binder and Simpson 2015), we can detect a range of interest in the bandwidth, and the impact of the window function on the final result is generally minor (Fig. S1). Finally, a comment on the errors for the 2pCF: The radial distribution function *c*_2*p*_ is generally computed without uncertainty bars attached. *c*_2*p*_ is indeed an average over many different measurements of the same histopathology slide. Therefore, despite being tiny because of working with many cells, the standard error in the mean could be worked out. Advanced bootstrap simulations found some attempts to account for this error in an astronomical context (Mo *et al* 1992). Another approach, different from what we have done, works on binned data, thus obtaining error bars as standard errors (Binder and Simpson 2015). This paper’s purpose remains to present a procedure whose nature of the data can be very different (from histopathology slides to MRI); therefore, the choice is presented in section 5.

### Supplement 3: Averages for ergodic non-equilibrium steady-state systems

The procedures to formalize the concept of “average” (used to describe the ROI in our histopathology slices) for non-equilibrium steady states in Sec.2.4 can be formalized with the help of the classical-quantum mechanic results (Landau and Lifshits 1974, Irving and Kirkwood 1950).

Following similar arguments common in the theory of stellar populations (Pasetto et al., 2019) for an arbitrary state variable *V* (e.g., cellular density in our histopathology slices), we consider the ROI as arbitrary non-equilibrium stationary systems. As introduced in the text, we assume each ROI locally represents the type of disease considered reasonably well. Therefore, the average non-equilibrium steady-state system then reads:

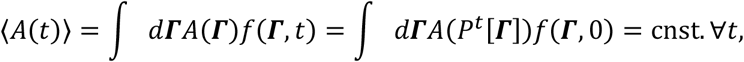

where it is easy to recognize the Schrodinger and the Heisenberg formalism to treat the distribution function of the cell distribution *f* in the first and second equality (defined in the phase-space ***Γ***). In the same way, as no a priori reason exists to prevent the cell the full access to the phase space (i.e., we assume to be far from any blood vessels, bones, etc., able to foliate/constraint the phase space), we can define the ergodic non-equilibrium stationary system as

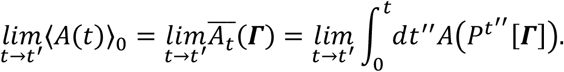

To investigate further the applicability of our argument in the interface ROI between stroma and tumor is beyond the goal of this work.

**Fig. S1.**
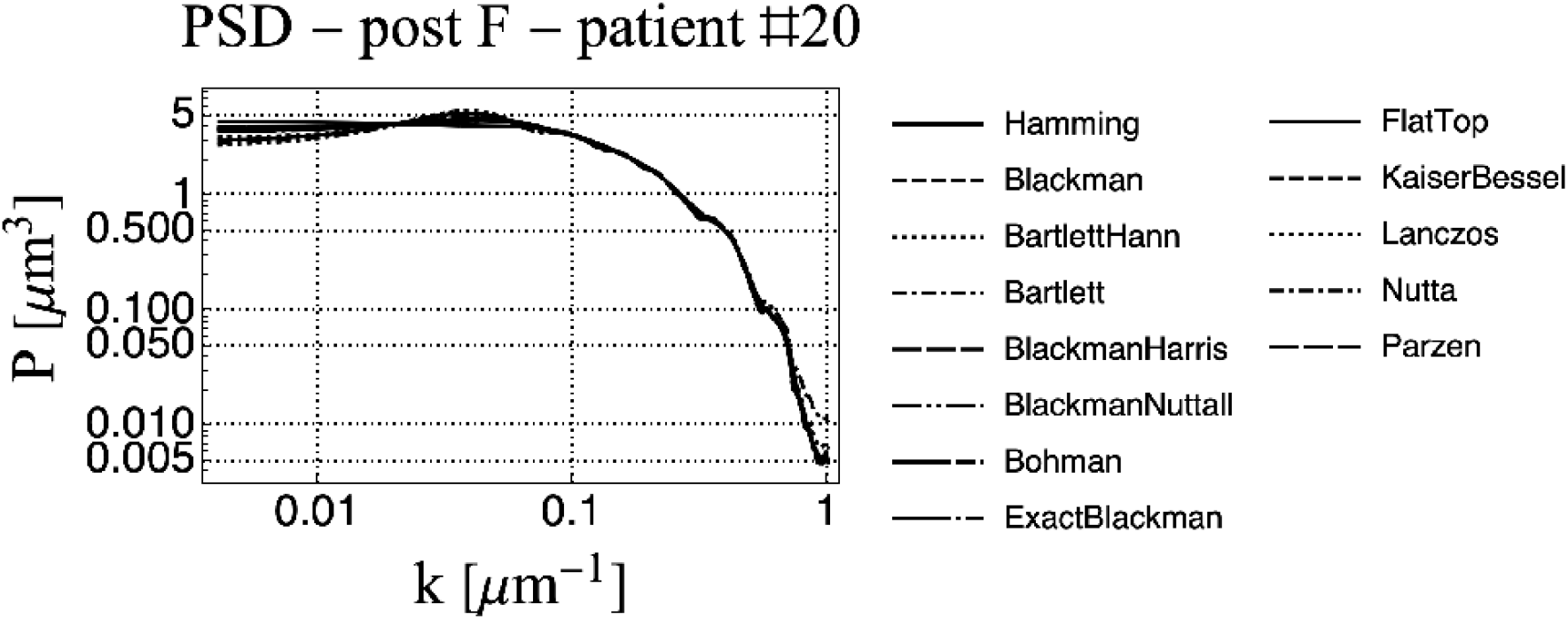
Different kernels impact. The impact of different kernels on the unnormalized PSD on patient #20 for the post-treatment area F.

**Fig. S2.**
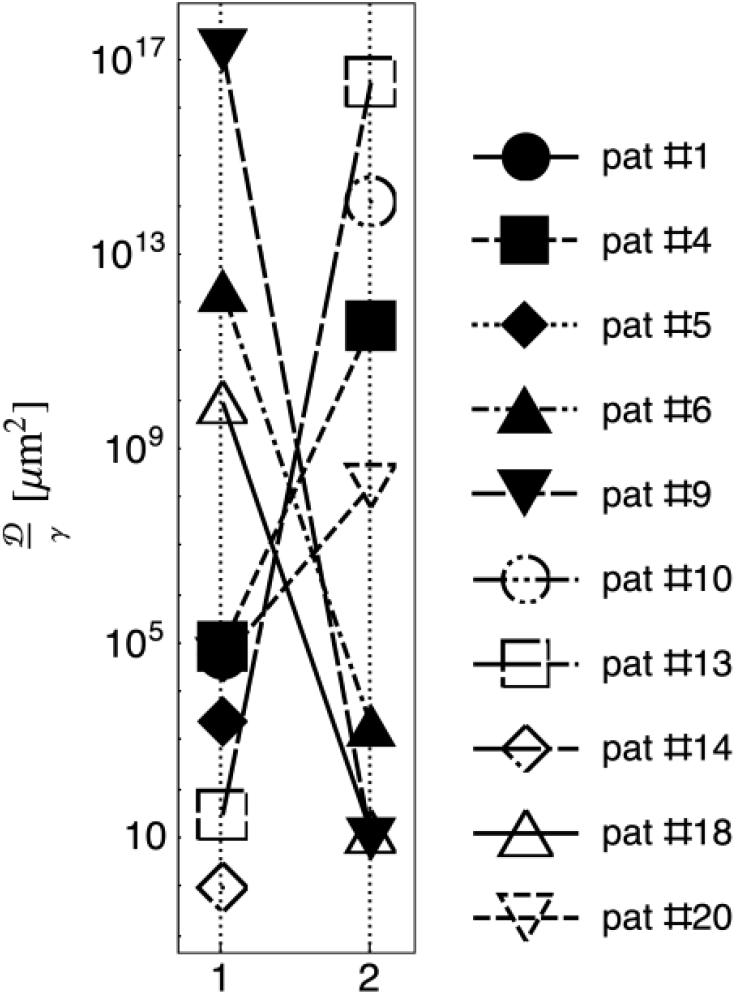
Diffusion over proliferation. Main text Fig 4c extended to include ten patients’ analyses.

## Footnotes

1 Note how the significant differences between npCf and cooccurrence (Haralick et al., 1973) are, aside from normalization factors, the pixeled/grid-free definition of the npCf and the possibility to capture both geometry and statistical nature of the images analyzed without grey-levels definition.

2 We note that the definition of the correlation function (or of the covariance function, or its relation with the Ripley’s K function) is not universal across the different disciplines and textbooks: the subtraction of the average density or any uncorrelated average product of two random variables is a matter of convention that, therefore we fix here with this definition.

